# A simple approach for accurate peptide quantification in MS-based proteomics

**DOI:** 10.1101/703397

**Authors:** Teresa Mendes Maia, An Staes, Kim Plasman, Jarne Pauwels, Katie Boucher, Andrea Argentini, Lennart Martens, Tony Montoye, Kris Gevaert, Francis Impens

## Abstract

Despite its growing popularity and use, bottom-up proteomics remains a complex analytical methodology. Its general workflow consists of three main steps: sample preparation, liquid chromatography coupled to tandem mass spectrometry (LC-MS/MS) and computational data analysis. Quality assessment of the different steps and components of this workflow is instrumental to identify technical flaws and to avoid loss of precious measurement time and sample material. However, assessment of the extent of sample losses along the sample preparation protocol, in particular after proteolytic digestion, is not yet routinely implemented because of the lack of an accurate and straightforward method to quantify peptides. Here, we report on the use of a microfluidic UV/visible spectrophotometer to quantify MS-ready peptides directly in MS loading solvent, consuming only 2 μl of sample. We determined the optimal peptide amount for LC-MS/MS analysis on a Q Exactive HF mass spectrometer using a dilution series of a commercial K562 cell digest. Careful evaluation of selected LC and MS parameters allowed us to define 3 μg as an optimal peptide amount to be injected on this particular LC-MS/MS system. Finally, using tryptic digests from human HEK293T cells, we showed that injecting equal peptide amounts, rather than approximated ones, results into less variable LC-MS/MS and protein quantification data. The obtained quality improvement together with easy implementation of the approach makes it possible to routinely quantify MS-ready peptides as a next step in daily proteomics quality control.

## Introduction

Mass spectrometry (MS)-based proteomics is a key technology in modern life sciences, with a wide scope of applications for protein research. Due to technological advances in liquid chromatography (LC) and MS instrumentation, current LC-MS/MS systems perform at high sensitivity, resolution and speed (Bache et al., 2018; D. Kelstrup et al., 2017; Espadas, Borràs, Chiva, & Sabidó, 2017; Meier et al., 2018). These improvements, coupled to robust bioinformatic pipelines now allow routine in-depth proteome measurements at high confidence, explaining the success of the major proteomics applications today: quantitative measurements of proteomes by so-called shotgun proteomics, identification of protein interaction partners by affinity purification MS (AP-MS), and mapping of the some of the most common post-translational modification (PTM) sites.

LC-MS/MS systems are inherently prone to fluctuations in performance. Whether an ion source that gets transiently unstable in the course of an analytical run, a mass analyzer with a drift in accuracy, or a chromatographic column whose stationary phase progressively deteriorates, the different components of an LC-MS/MS system vary during operation time (Tabb, 2013). Therefore, continuous monitoring is required to diagnose technical flaws and schedule maintenance interventions. For a very long time however, guidelines and standardized methods to assess the quality of MS-based proteomics data were lacking. Over the last decade, great progress has been made in proteomics quality control. On the one hand, various reference sample formulations have been designed and are now commercialized as standards to carry out QC analysis (Bittremieux, Tabb, et al., 2017)(Tabb, 2013). On the other hand, a number of metrics to assess LC-MS/MS performance have been developed (Bittremieux, Walzer, et al., 2017; Rudnick et al., 2010), in particular for standard workflows of bottom-up data-dependent analysis. Examples of QC metrics that inform about the liquid chromatography step are full-width at half-maximum (FWHM) of the eluting peptide peak and retention time drifts of eluting peptides, which monitor the integrity of the chromatographic column and the applied gradient. In turn, mass spectrometer QC can be done through examination of mass accuracy, dynamic range and ion injection times. Importantly, various software tools for MS-based proteomics quality control (QC) have been created (Bittremieux, Valkenborg, Martens, & Laukens, 2017), which allow extraction and visualization of QC metrics from analytical runs performed on standard samples. One such example is the cloud-based quality control system QCloud, which allows following instrument performance over time on a user-friendly web interface (Chiva et al., 2018).

Of utmost importance for the success of a proteomic experiment is the QC assessment at the sample preparation level, where the major sources of variation arise from (1) artificial and thus unwanted *in vitro* protein/peptide modifications, (2) incomplete protein digestion, (3) protein contamination introduced by users or present in reagents and (4) sample losses during sample transfer, enrichment steps, elution and reconstitution. While the extent at which the first three types of variation compromise the quality of an analytical run can be quantified (provided that the correct identification search settings are used), no standardized QC procedures exist that account for analyte losses after proteolytic digestion. Indeed, while the extracted amount of protein material from cell or tissue samples is typically determined by standard assays immediately upon lysis and/or protein extraction, or prior to digestion (Figure 1), no such established quantification procedures exist at the peptide level. Of particular note, even when additional modifications, enrichment steps or cleaning procedures take place after digestion, which will inevitably introduce additional sample losses and variability, peptide quantification before the actual LC-MS/MS analysis is generally not performed.

**Figure 1.**
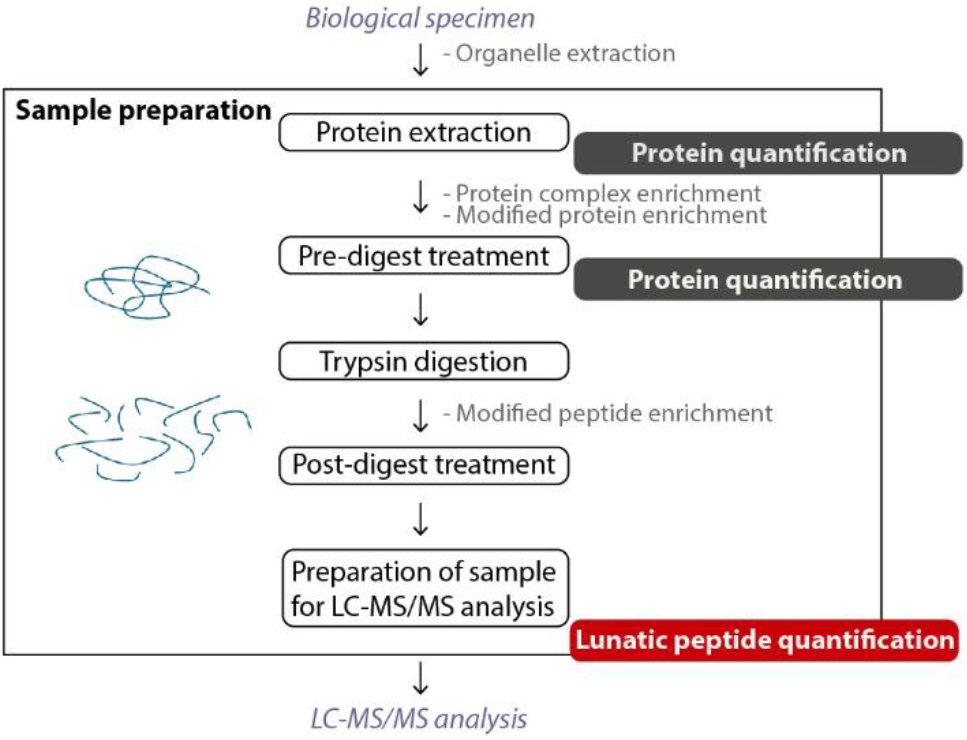
– Schematic outline of the sample preparation and quantification steps in a generic bottom-up proteomics experiment. Peptide concentration, which is usually estimated based on protein quantification before proteolytic digestion (dark gray), can be assessed right before LC-MS/MS using the Lunatic instrument (red). Optional steps of the protocol are written in gray.

LC-MS/MS runs carried out with insufficient amounts of peptides result in ion signals below the detection limit of the mass spectrometer, leading to low(er) numbers of peptide and protein identifications. In this respect, low signal to noise ratios of peptide signals also reduce the reliability of peptide quantification, as peak integration is more difficult for such low signals. Although analysis can be repeated by injecting more adequate sample amounts, this increases the overall analysis time and associated costs, and is not always possible for low quantity or low volume samples. Equally troublesome is the injection of excessive amounts of peptide material, termed column overloading. Such column saturation leads to peak tailing and retention time shifts (Johnson, Boyes, & Orlando, 2013; McCalley, 2003, 2010; Rieux, Lubda, Niederländer, Verpoorte, & Bischoff, 2006; Rieux, Sneekes, & Swart, n.d.), leading to poorer peptide separation and higher ionization competition, also reducing the number of identified and quantified peptides (Hinzke, Kouris, Hughes, Strous, & Kleiner, 2019). In some cases, this saturation can even lead to obstruction of the column, which causes back-pressure build-up, in its turn increasing the variability between repeated analyses of similar samples.

Efforts to standardize procedures and minimize sample losses during sample preparation led to the development of simplified uniform protocols, including protocols in which all pre-digestion steps, being lysis, protein reduction and alkylation are performed in a single volume (Batth et al., 2019; Kulak, Pichler, Paron, Nagaraj, & Mann, 2014; Moggridge, Sorensen, Morin, & Hughes, 2018). Solid phase extraction (SPE) to purify and concentrate peptides has also become common practice at the end of the sample preparation protocol. Furthermore, automated sample preparation workflows (Fu et al., 2018; Guryča et al., 2014) or robotized systems start to become available, which allow accurate liquid handling, temperature control and precise timing of processing steps (Kulak, Geyer, & Mann, 2017; Zhu et al., 2018). Some of these methods can be performed in miniaturized settings, which are advantageous when working with clinical or other low abundance samples where analyte amounts are scarce (Peng et al., 2016; Zhu et al., 2018).

Even when optimized sample preparation protocols are in place, straightforward QC assessment at the peptide level would be highly desirable for monitoring sample losses, leading to injecting optimal peptide amounts for LC-MS/MS analysis. In order for such a procedure to be aligned with a miniaturized, high-throughput mass spectrometry workflow, it should allow (1) batch-wise processing, (2) low sample consumption and (3) execution just before LC-MS/MS analysis in the loading solvent used. Here, we devised a procedure that complies with these requirements, based on peptide quantification in a microfluidics UV spectrophotometer (Lunatic, Unchained Labs). Consuming only 2 μl of sample, this instrument measures peptide concentrations as low as 31.25 ng/μl and returns the sample volume that should be injected for LC-MS/MS analysis of a desired peptide amount. Using this procedure, we determined the optimal amount of peptides to be injected for shotgun measurements on our LC-MS/MS setup. Finally, we demonstrated that the application of this procedure for control of sample loading improves LC-MS/MS and protein quantitation reproducibility.

## Methods

### Sample preparation

HEK293 cells (5 × 5×10^6^) were harvested, washed three times in PBS and resuspended in lysis buffer (8 M urea; 20 mM HEPES pH 8.0). Samples were sonicated by 10 pulses of 30 s at an amplitude of 20%, in a water bath kept at 10 ºC and centrifuged for 15 min at 20,000 g at room temperature to remove insoluble components. The protein concentration in the supernatants of each sample was measured using a Bradford assay (Bio-Rad) and equal protein amounts (500 μg each) were used for further analysis. Proteins were reduced with 5 mM DTT (Sigma-Aldrich) for 30 min at 55°C. Alkylation was performed by addition of 10 mM iodoacetamide (Sigma-Aldrich) for 15 min at room temperature in the dark. The samples were diluted with 20 mM HEPES pH 8.0 to a urea concentration of 4 M and proteins were pre-digested with 5 μg endoLysC (Wako, 129-02541) (1/100, w/w) for 4 h at 37 °C. All samples were further diluted with 20 mM HEPES pH 8.0 to a final urea concentration of 2 M and proteins were digested with 5 μg trypsin (V5111, Promega) (1/100, w/w) overnight at 37 °C. Peptides were then purified on a SampliQ SPE C18 cartridge (Agilent), vacuum dried and kept at −20 °C until further use.

### Peptide concentration measurements

All peptide concentration measurements were made on the Lunatic (Unchained labs) microfluidic spectrophotometer. Quantification of peptide mixtures was determined by the instrument’s “MS-peptide quant” tool, a software application that computes peptide concentration values based on UV-absorbance (280 nm) of aromatic amino acids.

### Linear dynamic range determination

The linear dynamic range for measurement of complex peptide mixtures on the Lunatic was determined on a group of eight solutions prepared from a 1.0 μg/μl stock of a commercial yeast protein digest (Promega, V7461), dissolved in loading solvent (0.1% TFA in acetonitrile/water, 2/98 (v/v)). Measurements of solutions in the 1.0 μg/μl to 10 ng/μl concentration range (used concentrations were 0.010, 0.025, 0.050, 0.100, 0.250, 0.500, 0.750, 1.000 μg/μl) were done in quadruplicate, from four yeast protein digest vials, and at least 2 readouts per solution were taken.

### Assessment of optimal loading amount

A commercially available K562 cell digest (Promega, V6951) was used to create a two-fold dilution series ranging from 0.013 to 1.0 μg/μl. Peptide amounts ranging from 0.06 μg up to 12 μg were loaded on a LC-MS/MS system. In parallel, the actual sample concentration values were assessed by Lunatic (see Figure 3a). The same experiment was repeated using a new K562 cell vial to prepare the dilution series and a different analytical column.

Assessment of LC-MS/MS analysis quality under controlled sample loading conditions

Five HEK293 protein digest samples were resuspended in 30 μl MS loading solvent (0.1% TFA in acetonitrile/water, 2/98 (v/v)) and measured on Lunatic. Two sets of LC-MS/MS analytical runs followed. In the first one, 3.0 μg of peptide mixture from each sample was loaded onto the LC-MS/MS system, while in the second one the same sample volume was used for all five injections, resulting in peptide amounts ranging from 2.50 to 3.55 μg (detailed in the Results section).

### LC-MS/MS analysis

From each sample, 5 μl was introduced into an LC-MS/MS system through an Ultimate 3000 RSLC nano LC (Thermo Fisher Scientific, Bremen, Germany) in-line connected to a Q Exactive HF mass spectrometer (Thermo Fisher Scientific). The sample mixture was first loaded on a trapping column (made in-house, 100 μm internal diameter (I.D.) × 20 mm, 5 μm beads C18 Reprosil-HD, Dr. Maisch, Ammerbuch-Entringen, Germany). After flushing from the trapping column, the sample was loaded on an analytical column (made in-house, 75 μm I.D. × 400 mm, 1.9 μm beads C18 Reprosil-HD, Dr. Maisch) packed in the needle (pulled in-house). Peptides were loaded with loading solvent (0.1% TFA in acetonitrile/water, 2/98 (v/v)) and eluted with a non-linear 150 min gradient of 2-56% solvent B (0.1% formic acid in water/acetonitrile, 20/80 (v/v)) at a flow rate of 250 nL/min. This was followed by a 15 min wash reaching 99% solvent B and re-equilibration with solvent A (0.1% formic acid). The column temperature was kept constant at 50 °C (CoControl 3.3.05, Sonation).

The mass spectrometer was operated in data-dependent, positive ionization mode, automatically switching between MS and MS/MS acquisition for the 16 most abundant peaks in a given MS spectrum. The source voltage was set to 3.0 kV and the capillary temperature was 250 °C. Full scan MS spectra (375-1,500 m/z, AGC target 3 × 10^6^ ions, maximum ion injection time of 45 ms) were acquired at a resolution of 60,000 (at 200 m/z) in the orbitrap analyzer, followed by up to 16 tandem MS scans (resolution 15,000 at 200 m/z) of the most intense ions fulfilling predefined selection criteria (AGC target 1 × 10^5^ ions, maximum ion injection time of 60 ms, isolation window of 1.5 m/z, fixed first mass of 145 m/z, spectrum data type: centroid, underfill ratio 2%, intensity threshold 1.3 × 10^4^, exclusion of unassigned, singly and >7 charged precursors, peptide match preferred, exclude isotopes on, dynamic exclusion time of 12 s). The HCD collision energy was set to 28% of the normalized collision energy and the polydimethylcyclosiloxane background ion at 445.12002 Da was used for internal calibration (lock mass).

### Data analysis

Data analysis was performed with the MaxQuant software (version 1.6.1.0) (Cox & Mann, 2008). The Andromeda search engine was used with default settings, including PSM, peptide and protein false discovery rate set at 1% and match between runs disabled. Spectra were searched against the UniProt reference proteome release 2018_01 (UP000005640_9606, containing 21007 human protein entries) with a mass tolerance for precursor and fragment ions of 4.5 and 20 ppm, respectively. Oxidation of methionine residues and acetylation of protein N-termini were defined as variable modifications, while carbamidomethylation of cysteine residues was set as a fixed modification. Proteins with at least one unique or razor peptide were retained, then quantified by the MaxLFQ algorithm integrated in the MaxQuant software (Cox et al., 2014). A minimum ratio count of two unique or razor peptides was required for quantification. Further data analysis was performed with the Perseus software (version 1.6.1.3) after loading the protein groups file from MaxQuant. The number of protein identifications was given by the number of protein groups obtained after filtering out protein groups only identified by site, reverse database hits and potential contaminants. The number of peptide identifications was obtained after exclusion of reverse database hits and potential contaminants. Quantified proteins were the subset of identified proteins with positive LFQ intensities. Heat maps were prepared from matrices of log2 transformed LFQ protein expression values, after filtering for at least three valid values over all samples. Proteins were displayed according to the order of groups defined by hierarchical clustering, using Euclidean distances and average linkage settings.

The chromatographic parameters signal to noise (S/N) ratio and peak area were obtained from the moFF algorithm (Argentini et al., 2016), run on each analytical run. Peak areas were approximated by the area of a triangle of base equal to the peak width (in minutes) and height given by the peak’s apex intensity. MS2 injection time was retrieved from MaxQuant’s MS/MS scans table. This analysis was done on peptide-to-spectrum matches from features detected in every analytical run from each experiment. Peptides were partitioned according to hydrophobicity the following way: hydrophilic peptides were those eluting until 50 min of the LC gradient (1.6-12.3 % acetonitrile), intermediate hydrophobic peptides eluted between 50 and 110 min (12.3-26.6 % acetonitrile) and hydrophobic peptides eluted after 110 min (26.6-79.2 % acetonitrile) (Figure S1).

The mass spectrometry proteomics data have been deposited to the ProteomeXchange Consortium (http://proteomecentral.proteomexchange.org) via the PRIDE partner repository with the identifier 10.6019/PXD014524 (username; password: 2RDeDBQ4).

## Results and Discussion

The introduction of a peptide purification step prior to LC-MS/MS analysis (Peng et al., 2016) in routine in proteomics laboratories opened an opportunity to accurately determine peptide concentrations. Such a peptide purification step, typically based on Solid Phase Extraction (SPE), removes buffer components (like chaotropes, certain detergents, reducing/alkylating reagents and trypsin) that interfere with LC-MS/MS analysis. As these components often absorb UV light, for long time they prevented routine, fast and accurate peptide quantification, often leading to the analysis of suboptimal peptide amounts by LC-MS/MS. The high purity of peptides obtained after SPE led us to explore the possibility to quantify MS-ready peptide samples using a Lunatic spectrophotometer from Unchained Labs.

A simple application for accurate quantification of MS-ready peptide samples

The Lunatic instrument is a microfluidic spectrophotometer that performs DNA, RNA, protein and peptide quantifications in batch, consuming only 2 μl of sample. Determination of peptide concentrations occurs accurately by a proprietary software application that, from an acquired UV-visible absorbance spectrum, discriminates the signal coming from the UV light absorbing amino acids from those of usual contaminants (e.g. nucleic acids, cofactors). The “MS-peptide Quant” application, available since September 2017 on Lunatic instruments, quantifies peptides by taking into account the extinction coefficients of tryptophan and tyrosine at 280 nm, as well as estimated values for both their fractions in a protein digest and their proportion in mass relative to the whole protein content of a sample.

The ability to quantify complex peptide mixtures was verified using dilution series of commercial protein digests from yeast, ranging in concentration between 10 ng/μl and 1 μg/μl, based on the theoretical peptide quantity as reported by the vendor. Comparison between the theoretical concentration values and the measured ones in four independent replicate dilutions showed a high linear correlation (R2 >0.98) and a slope very close to 1 (t-test p-value = 0.19), demonstrating the instrument’s high accuracy (Figure 2). It must be noted that peptide solutions with concentrations below 50 ng/μl gave absorbance measurements lower than 0.03, the lower absorbance limit at which analyte concentration can be accurately assessed according to the Lunatic specifications. Excluding measurements from such samples, the coefficient of determination (R2) between the theoretical and observed peptide concentrations was above 0.98. Therefore, it was concluded that solutions of peptides with concentrations between 50 ng/μl and 1 μg/μl are within the dynamic range of the instrument and can thus be accurately measured. This concentration range is well suited for MS-ready samples, for which LC-MS/MS experiments typically consume peptide quantities between 100 ng and 5 μg, contained in a volume of a few μl.

**Figure 2.**
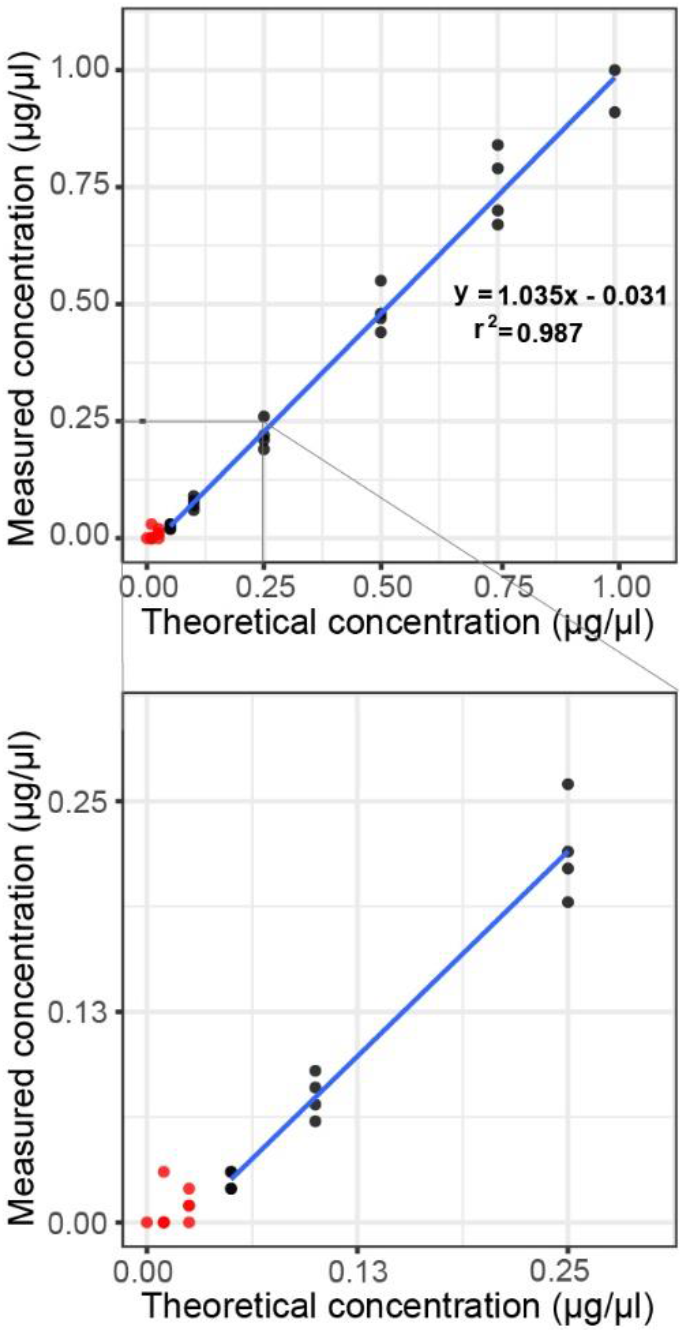
– Dynamic quantification range of the Lunatic spectrophotometer for complex peptide mixtures. Yeast peptide digest solutions with concentrations ranging from 10 ng/μl to 1.0 μg/μl (black dots) can be accurately measured on the Lunatic, as shown by a strong correlation between the theoretical and measured peptide concentration values. Below 50 ng/μl (red dots), peptide measurements are no longer reliable, as absorbance values reach the lower limit of detection of the spectrophotometer. Dots represent measurements of four independent replicate dilutions.

In summary, together with its accuracy for measurement of peptide solutions at commonly used concentration ranges, peptide quantitation with the Lunatic instrument presents a unique set of characteristics that make it particularly attractive over other quantification methods for MS-ready bottom-up proteomics samples (e.g. Bradford, Lowry, BCA, Pierce 660 nm, tryptophan fluorescence (Wiśniewski & Gaugaz, 2015)). It is a quick and easy method that can be performed in batch. It only implies a very low, 2 μl, sample consumption. It is based on UV-light absorbance at 280 nm, an intrinsic property of aromatic amino acids, thus requiring no calibration curve, unlike colorimetric reaction-based assays like Bradford or BCA.

Determination of an optimal peptide amount for LC-MS/MS analysis

The ideal peptide amount to use for LC-MS/MS in order to have a maximum throughput and identification rate, while keeping a good analyte separation, will largely depend on the characteristics of the LC-MS/MS system, namely on its chromatographic column loading capacity. If the exact sample concentration would be known, it would be possible to always work with optimal sample amounts, and thus consistently get high quality LC-MS/MS runs. Therefore, taking advantage of peptide quantification by the Lunatic instrument, we set out to determine the most suitable sample loading for an LC-MS/MS setup routinely used in our laboratory: a 40 cm LC column packed in the needle with 1.9 μm C18 beads coupled to a Q Exactive HF mass spectrometer.

Starting from a commercial tryptic digest of a K562 cell lysate, we prepared 10 different samples with peptide concentrations ranging from 0.013 μg/μl to 2.4 μg/μl. Five microliter of these samples, containing peptide amounts ranging from 0.06 to 12 μg, was injected for LC-MS/MS analysis using a 2.5 hour gradient. The experiment was repeated to generate data for two independent series of replicates analyzed on two different LC columns (Figure 3, Figure S2).

**Figure 3.**
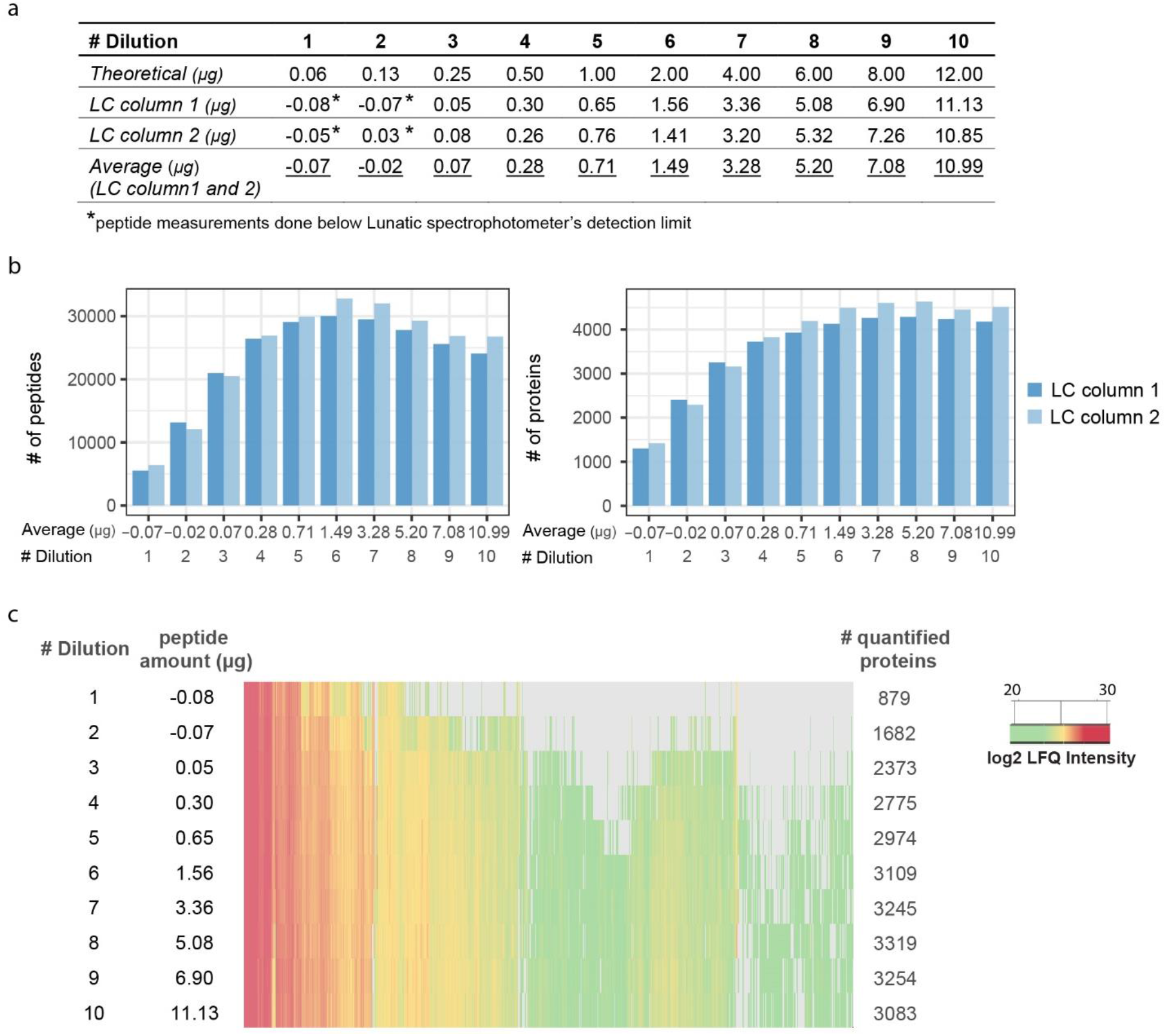
Identified and quantified peptide and protein numbers upon LC-MS/MS analysis of 10 different amounts of a trypsin digested K562 cell extract. (a) Theoretical and measured peptide amounts for two independent experiments. In both experiments, all measured values were lower than predicted, pointing to peptide losses during sample solution preparation. (b) Bar plots indicating the numbers of identified peptide and proteins per analytical run. Colors represent the results of two replicate series analyzed on two different LC columns. (c) Representative heatmap visualizing the intensities of quantified proteins in each of the 10 analytical runs measured using LC column 1. Missing values are shown in gray. Based on numbers of identified and quantified proteins a rather broad window ranging from 1.49 to 5.20 μg of peptides was found as an optimal injection amount.

All peptide concentration measurements were lower than predicted (Figure 3a). Likely, these deviations arose mostly from peptide losses during peptide resuspension of the K562 digest. Additional losses during the preparation of serially diluted samples were minor, judging from the ratios between consecutive dilutions, which were very close to expected (R^2^ >0.99). We also noticed that peptide quantification of # Dilution 1 and 2, having theoretical concentrations of 0.013 and 0.025 μg/μl, returned negative values. These clearly inaccurate measurements were performed below the limit of detection of the spectrophotometer.

To find an optimal peptide amount to inject on our LC-MS/MS setup, we evaluated the number of identified and quantified proteins. To this end, the LC-MS/MS data were searched with MaxQuant (Cox & Mann, 2008) and identified proteins were quantified by the MaxLFQ algorithm (Cox et al., 2014) (Figure 3b-c, Figure S2, Table S1). We observed that the number of identified proteins increased with the injected peptide amount until it started to level off at an average amount of 3.28 to 5.20 μg on injected peptides. As for the numbers of identified peptides, these reached a maximum between 1.49 to 3.28 μg. This suggested that column overloading occurred for peptide amounts above 3.28 μg, leading to a marked decrease in the number of identified peptides (18 to 20% reduction in numbers of identified peptides for the 10.99 μg samples). Although these peptide losses did not readily translate into a lower number of identified proteins, they did have an impact on the number of quantified proteins. Indeed, injection of an average amount of 3.28 to 5.20 μg of peptides resulted in the highest number of quantified proteins (Figure 3c, Figure S2, Table S1). Taken together, based on numbers of identified and quantified proteins, a rather broad window ranging from approximately 1.49 to 5.20 μg of peptides was found as an optimal peptide injection amount for our LC-MS/MS setup.

In an attempt to further narrow down this window, we decided to evaluate selected LC and MS parameters. The efficient time consumption of a Q-Exactive HF mass spectrometer is, among other things, reflected by the ion injection time. The closer to ideal the amount of material presented to the mass spectrometer is, the lower the injection time will be. Figure 4a (left panel) shows that the average MS2 ion injection time decreases with increasing peptide amount, leveling off and reaching an optimal minimum around an average amount of 3.28 μg of injected peptides. We also assessed the signal-to-noise (S/N) ratio of chromatographic peaks and peak areas, two LC parameters that were extracted for all identified peptides using the moFF algorithm (Argentini et al., 2016). Figure 4a (right panel) shows that the average S/N ratio increases with increasing peptide amounts and again levels off at an average amount of about 3.28 μg of injected peptides. Analysis of the median peak area of all identified peptides showed a maximum between 5.20 and 7.08 μg for the two replicate experiments (Figure 4a (middle)). When overloading a reversed phase C18 column, one expects peak tailing, with a concomitant increase of the average peak area due to repulsion effects of same-charge ions (Johnson et al., 2013; McCalley, 2003, 2010). The observed leveling off and even decrease of the peak area at higher loads though contradicted this expectation. To explore this further, we split peptides according to their retention time interval into three parts: early eluting peptides (hydrophilic, n=126), late eluting peptides (hydrophobic, n=178) and the ones eluting in between (mid eluting/intermediate hydrophobic peptides, n=2304) (Figure S1). This classification revealed an interesting trend especially for the hydrophilic peptides, consisting of a considerable decline in peak area starting from around 2-3 μg for both replicate series (Figure 4b), pointing to a loss of these peptides.

**Figure 4.**
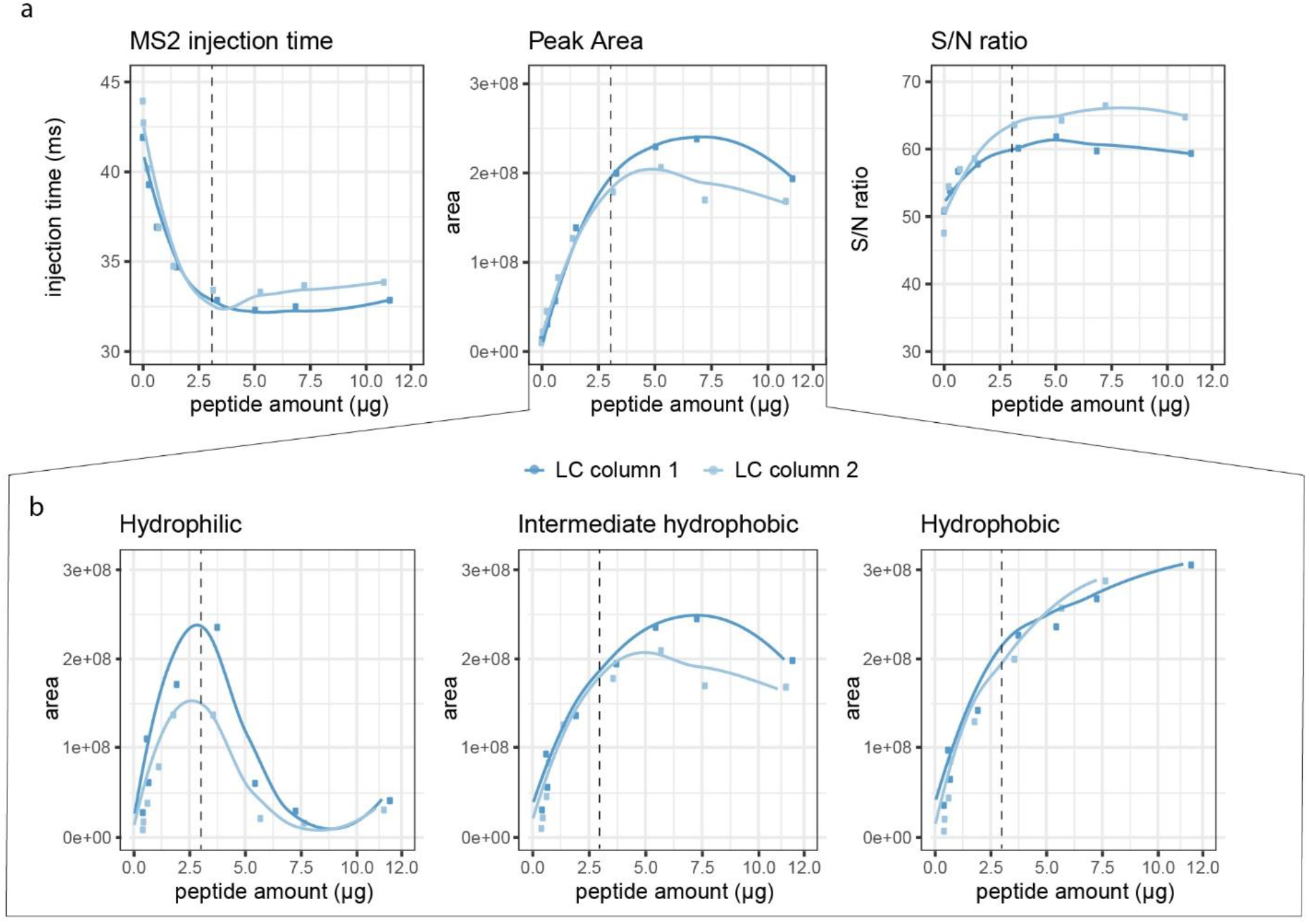
Liquid chromatography and mass spectrometry metrics upon LC-MS/MS analysis of 10 different amounts of a trypsin digested K562 cell extract. (a) Scatter plots of average MS2 ion injection time, median peak area and median signal-to-noise ratio of identified peptides in each analytical run. (b) Scatter plots of the median peak area of hydrophilic, intermediate hydrophobic and hydrophobic identified peptides. Colors represent the results of two replicate series analyzed on two different LC columns. Loess curve fitting was used to generate the connecting lines. Based on the loss of hydrophilic peptides 3 μg was defined as optimal peptide injection amount on this particular LC-MS/MS setup (dashed line).

This loss is most likely related to the use of a trapping column in our LC-MS/MS setup, whose capacity appears not to allow retention of the most hydrophilic peptides for higher sample amounts. In contrast, the hydrophobic peptides, having a higher affinity for the stationary phase, do not elute from the trapping column during the washing step that precedes the analytical column chromatography. As a consequence, the more peptides were loaded, the more the peak area of hydrophobic peptides would increase, as shown in Figure 4b (right panel). The peptides belonging to the middle region do not have a pronounced affinity for the stationary phase, and will therefore have a mixed behavior (Figure 4b, middle panel). The disadvantage of losing hydrophilic peptides does not overrule the advantages of using a trapping column though, if void volumes are avoided (Mitulović et al., 2003). Therefore, based on the loss of hydrophilic peptides for amounts above 2.86 μg and 2.63 μg for replicate series 1 and 2, respectively (peptide amounts that correspond to the peak maxima in the hydrophilic peptides peak area plot), together with the stabilization of the other parameters also around these sample amounts, we now routinely use 3.0 μg of peptides as an optimal injection amount on this particular LC-MS/MS setup.

By running a series of LC-MS/MS analysis, covering a range of known peptide sample amounts, and performing a follow-up evaluation of the numbers of identified and quantified features, as well as of a few LC and MS parameters, we were able to identify 3 μg as an optimal amount of injected peptides for our LC-MS/MS setup. This procedure could be applied to any other system and as such be of great use in proteomics laboratories. Indeed, knowledge about the most adequate sample amounts to use is currently not known for most systems. As a workaround to deal with this lack of information, it is common to, either inject different concentrations of the same sample, or else to perform short off-line analytical runs to fine-tune sample loading for a given sample (Pynn, Christopher Samonig, Krssakova, Mechtler, Decrop, & Swart, n.d.). However, these strategies are time and cost ineffective and often consume higher amounts of precious sample material than the simple UV-based procedure described here.

Control over peptide amount injections effectively stabilizes LC-MS/MS and protein quantitation reproducibility

During sample preparation, it is common to make the protein amounts for all samples of a batch even before proceeding to the digestion step. This will generate very equivalent MS samples that can then be injected for LC-MS/MS using equal volumes. We decided to evaluate if, in comparison to that approach, performing peptide quantification prior to MS and concomitantly adjusting injection volumes to account for small differences in samples concentrations would have an impact on the reproducibility of LC-MS/MS and, in particular, of quantitation data.

To this end, we analyzed five tryptic digests of HEK293T cells, prepared in parallel starting from five different cell pellets. On a first set of analytical runs, we corrected for the peptide amounts loaded based on the Lunatic concentration, having injected exactly 3.0 μg of peptide mixture for all five replicates. In contrast, on a second set of runs of the same samples, we injected a fixed volume, corresponding to the average used for the first experiment. After calculating the peptide loadings for this second experiment, we realized there was a difference of 1 μg between the lowest and highest concentration sample (2.5 μg and 3.55 μg, respectively). These differences are quite substantial and show the importance of accurate peptide quantitation of MS-ready samples. Even with automation of sample preparation, which should alleviate heterogeneity between samples, peptide quantitation could always be kept as a quality check step.

For both sets, the previously used MS and LC parameters were examined (Figure 5a). Strikingly, while all MS injection time, peak area and S/N noise ratio values were close to each other in the experiment with the “corrected” sample amounts, these showed a much higher spread in the second experiment where each analytical run had a different sample amount. Indeed, the standard deviation for MS2 injection time, peak area and S/N ratio had a 10-fold, a 3-fold and a 2-fold increase, respectively, for the ‘non-corrected’ experiment relative to the ‘corrected’ experiment (Figure 5a).

**Figure 5.**
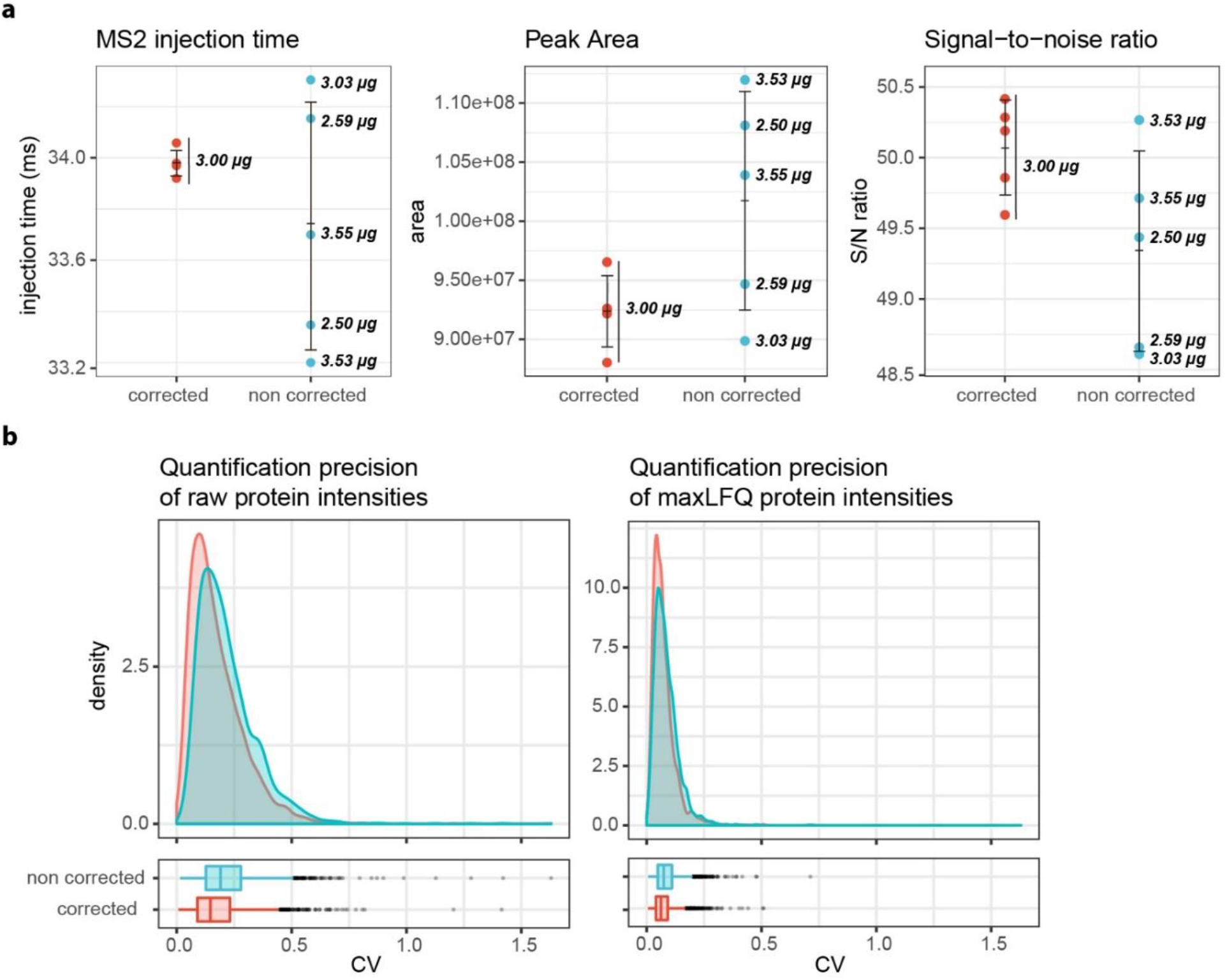
– Comparison of LC-MS/MS and protein quantification reproducibility for two sets of analytical runs on trypsin digested Hek293T cell extracts. (a) Plots of the average MS2 injection time, median peak area and median signal-to-noise ratio values are shown for the identified peptides in each of five replicates from the ‘corrected’ and ‘non corrected’ groups of samples. For each data point, the exact injected peptide amount based on Lunatic quantification is shown on the right. Vertical bars represent mean ± standard deviation. (b) Density plots and boxplots showing the distribution of the coefficients of variation (CV) of raw intensities (left) and (normalized) maxLFQ intensities (right) for all proteins quantified in the 10 analytical runs (n=2677). For both raw and normalized protein intensities, a shift to higher CV values was found for the ‘non-corrected’ samples (Mann-Whitney U test pvalue < 2.2 x10^−16^ for the shift in both raw and normalized distributions).

Finally, we checked if this higher analytical reproducibility also resulted in a more precise protein quantification. To this end, we looked at the distribution of the coefficients of variation (CV) of raw protein intensity values from corrected replicates versus that from non-corrected replicates. In addition, we compared the CV distributions for LFQ protein intensity values obtained after normalization by the MaxLFQ algorithm. As expected, CV values for the non-normalized intensities were significantly lower for the ‘corrected’ experiment (Figure 5b, Mann-Whitney U test p-value < 2.2 ×10^−16^). However, a shift towards lower CV values was also observed for the normalized LFQ intensities of the “corrected” experiment (Mann-Whitney U test p-value < 2.2 ×10^−16^), indicating that computational normalization cannot fully compensate for differences in injection amounts. Such reduction in the dispersion of protein quantification estimates can potentially improve the sensitivity for detecting peptide and protein regulation. Together, these data show that working consistently with equal peptide amounts results in increased reproducibility LCMS/MS and protein quantification data, even in combination with normalization of protein intensities during data analysis.

## Conclusion

In the present work, we describe a simple procedure to quantify MS-ready peptides using the Lunatic microfluidic UV-visible spectrophotometer from Unchained Labs. The Lunatic instrument, with its integrated peptide quantification software application, constitutes an accurate and easyto-use device for the implementation of peptide quantification in proteomics laboratories and facilities. Through careful control of the injected peptide amount on a commonly used LC-MS/MS setup, we show how this procedure can provide a constant quality boost to MS-based proteome analyses. Application of this methodology on a routine basis can drastically reduce time and sample losses by avoiding re-runs after injection of suboptimal peptide amounts.

## Supporting information

SFigure 1

SFigure 2

## Supporting Information

Figure S1 – Schematic plot showing the percent solvent B gradient and total ion current along retention time from a representative LC-MS/MS run of the sample amount optimization experiment. Retention time points used for peptide partioning are also shown.

Figure S2 – Heatmap visualizing the intensities of quantified proteins in each of the 10 analytical runs measured using LC column 2. Missing values are shown in gray.

Table S1 – Numbers of identified and quantified peptide and protein groups for the two replicate series of K562 analytical runs. This material is available free of charge via the Internet at http://pubs.acs.org.

## AUTHOR INFORMATION

### Author Contributions

K.P., T.M., F.I., K.G., A.S and J.P. conceptualized the study and designed the experiments. K.P. and T.M.M. prepared the yeast and K562 cell peptide dilution samples. K.B prepared the Hek293T proteome digests. J.P. performed the LCMS/MS measurements. T.M., A.S, A.A and J.P. analyzed the data. T.M wrote the manuscript with input from F.I., K.G., L.M. All authors revised the final manuscript.

### Funding Sources

The authors would like to acknowledge the VIB Tech Watch Fund for supporting early access to innovative technology platforms. L.M., K.G. and F.I. acknowledge support by the EPIC-XS consortium and F.I. acknowledges support by FWO-SBO Project S006617N.

### Notes

The authors declare no competing financial interest.

## ACKNOWLEDGMENT

The authors thank Lisa Adamiak and Dina Finan from Unchained Labs for technical advice and sharing information about the Lunatic instrument specifications.

## ABBREVIATIONS

LC-MS/MS: liquid chromatography-tandem mass spectrometry
LFQ: label free quantification

---

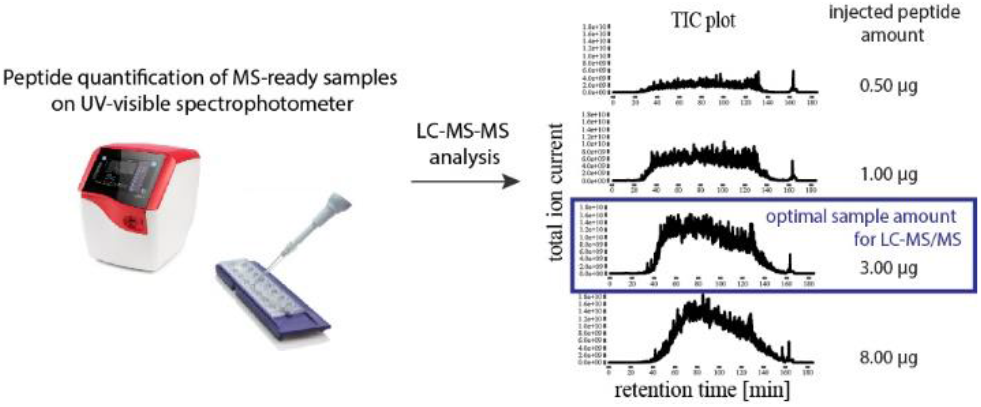

