## Supplementary figures and images for "A simple approach for accurate peptide quantification in MS-based proteomics"

### SFigure 1

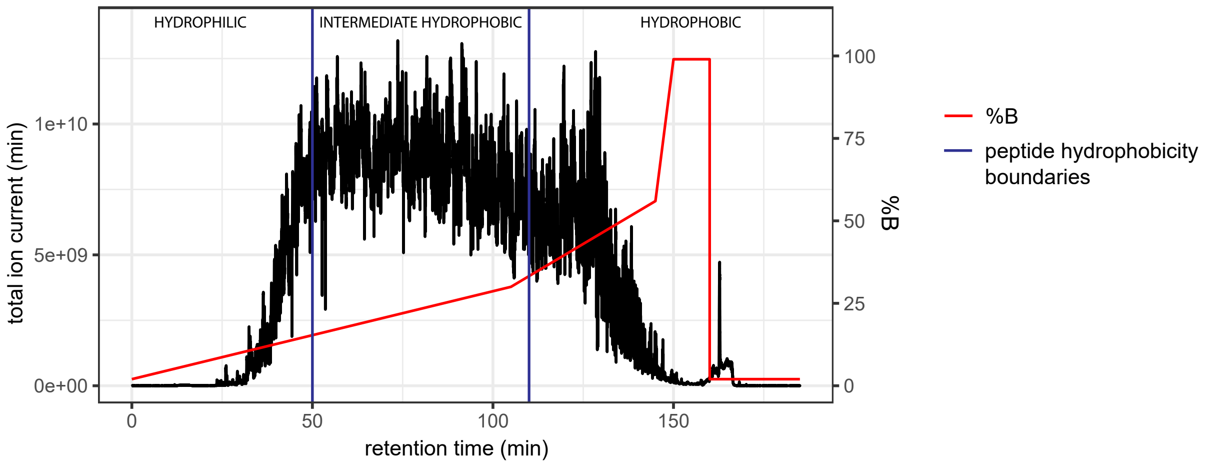

### SFigure 2

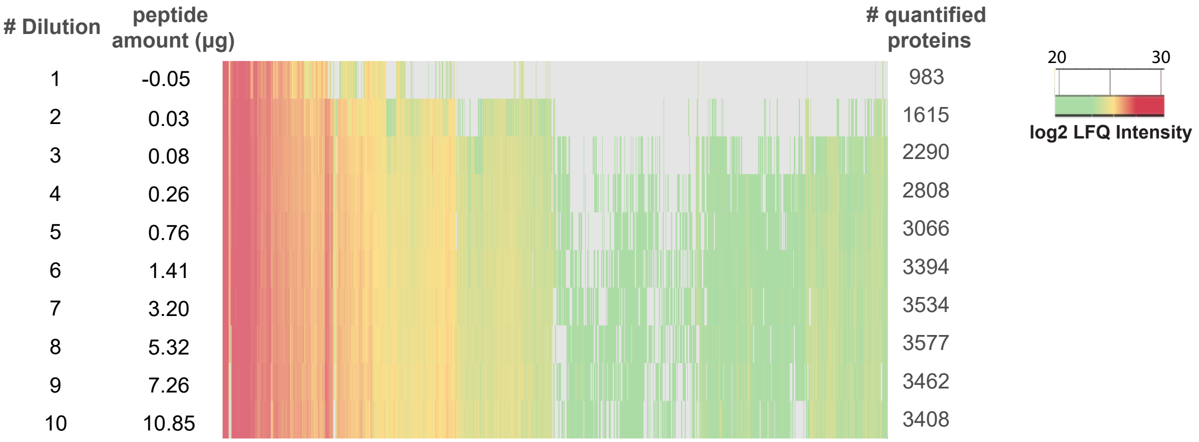
